# Synchronized Chromatin Organization Serves as Potent Biomarker in Anaesthesia-based Plant Consciousness

**DOI:** 10.1101/2024.09.27.615456

**Authors:** Shilpa Chandra, Bodhidipra Mukherjee, Abdul Salam, Farhan Anjum, Chayan Kanti Nandi, Laxmidhar Behera

**Affiliations:** Centre for Indian Knowledge System and Mental Health Applications, Indian Institute of Technology Mandi, HP-175005; School of Biosciences and Bioengineering, Indian Institute of Technology Mandi, HP-175005; School of Chemical Science, Indian Institute of Technology Mandi, HP-175005; School of Computer Sciences and Electrical Engineering, Indian Institute of Technology Mandi, HP-175005

**Keywords:** Anaesthesia, Consciousness, Cellular organelle, Plant, Chromatin, Biomarker

## Abstract

Anaesthesia has been used for centuries for medical purposes. With the application of anaesthesia, organisms lose their conscious awareness. It provides a temporary loss of sensation, which enables painless performance during surgery. However, the cellular mechanisms underlying the effects of anaesthesia are not clearly understood. It has been proposed that plant root function is analogous to the human brain. Here, using super-resolution imaging technique, we explored an organelle-level understanding of the effect of anaesthesia on plant roots and the stem connecting to the root. Our results showed that the nuclei organized themselves in an orchestrated manner upon treatment with both local and general anaesthesia without damaging their structure. Euchromatin within the nucleus was found to be reorganized in the nuclear periphery, and this process was found to be independent of ATP. In contrast, mitochondria, microtubules, endocytic vesicles, and chloroplasts, which are other important organelles in plant cells, were highly altered or damaged under the same experimental conditions. Eventually, the cellular homeostasis again maintained and process is reversible upon the removal of anaesthesia. Our results suggest that such orchestrated chromatin organization without disturbing the overall structure of the nucleus could be used as a potent biomarker for conscious awareness in plants.

**Graphical Abstract:** 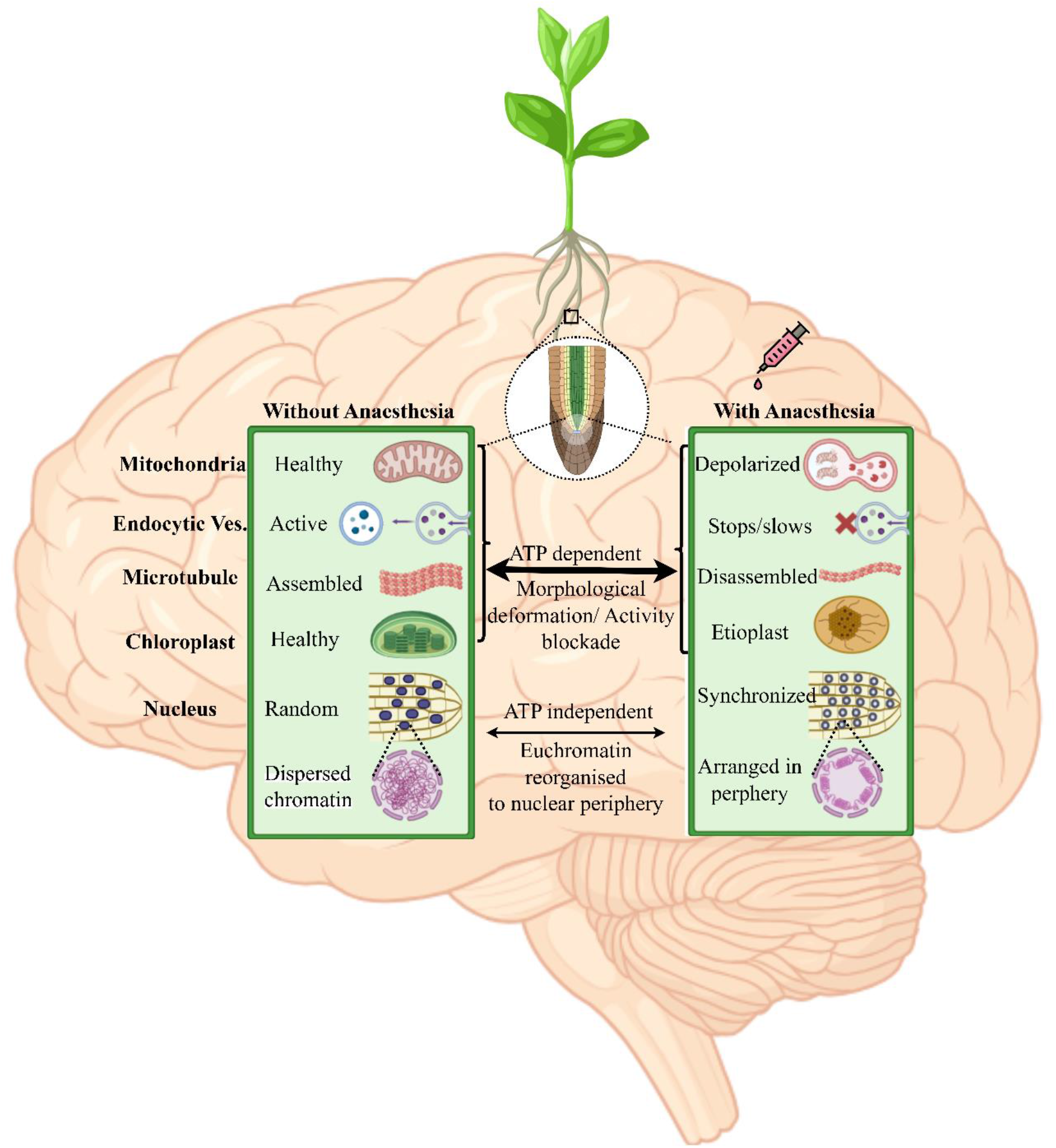

## Introduction

Anaesthesia has been employed in medical procedures for centuries since its effects have been inadvertently discovered. Despite extensive efforts to provide a scientific rationale for the impact of these substances, the question of why a wide array of chemical compounds from different categories can induce the shared effect of unconsciousness in both animals and plants remains ambiguous^1^. Remarkably, anaesthesia treatment does not eliminate the underlying state of conscious awareness; otherwise, the organism dies ^2^. All organisms share fundamental traits that are universal to all living beings, such as sensory systems, ability to act, cognitive abilities, behavior, and capacity to adapt. Anaesthesia works by producing a temporary state of loss of sensation, particularly noxious stimuli, and conscious awareness of the environment without causing harm to the individual or their biological cells. Rather, it only delays the transmission of signals, making one feel no pain, and causing unconsciousness during medical procedures ^3–7^.

An essential part of environmental consciousness is the ability to perceive and analyze information from outside sources ^8^. Animals typically require sophisticated mental processes alongside their highly evolved sensory systems (sight, sound, touch, taste, and smell) ^9^. While animals have central nervous systems, plants do not have nervous systems; they sense their surroundings and react accordingly. Their exceptionally refined sensory systems can detect a wide variety of environmental cues including chemicals, light, gravity, and touch ^10^. Many studies have suggested that plant roots function analogously to the brain by coordinating and integrating responses to environmental stimuli^11^. The root apex, particularly the root cap, contains cells that perceive and process information, facilitating complex behaviors such as growth direction and resource allocation. This concept, known as the “root brain” hypothesis, proposes that the roots exhibit a form of decentralized intelligence, wherein the root meristem acts as a command center, processing sensory inputs, and guiding adaptive responses^12^. This capability is underpinned by a sophisticated signaling network involving hormonal gradients, electrical signals, and gene expression changes, allowing plants to navigate and optimize their growth in a dynamic environment ^13,14^. Recent studies have shown that plants produce sound when they are stressed. They can communicate with each other using volatile organic compounds, electrical signaling, and common mycorrhizal networks between plants and a host of other organisms such as soil microbes, plants of the same or other species, animals, insects, and fungi ^15,16^.

Reports have suggested that the anaesthetic effect in plants, such as Mimosa pudica L., was unresponsive in closing leaves upon contact with the anaesthetic compound diethyl ether ^17^. Following exposure to anaesthetics, mimosa leaves, pea tendrils, Venus flytraps, and sundew traps all lost their ability to move autonomously and in response to touch. This phenomenon was explained by the loss of action potentials during anaesthesia treatment. ^14^. This observation led researchers to conclude that plants and animals share a common biological essence that is disrupted by anaesthesia^18–20^. There are few studies on imaging of plant organelles dynamics under anaesthetic conditions. Imaging of Arabidopsis cells has shown that anaesthesia induces reactive oxygen species (ROS) production and disrupts endocytic vesicle recycling^14^. Additionally, dormancy termination and chlorophyll biosynthesis were significantly hindered during anaesthesia^14,21^.

Despite the above studies, the organelle level underpinning and the specific mechanisms by which anaesthesia bearing different chemical structures causes similar types of unconsciousness are still unclear. Studied by Stuart Hameroff and Roger Penrose formulated the Orchestrated Objective Reduction (Orch-OR) theory, proposing that consciousness arises from quantum events occurring in microtubules present in brain neurons ^22^. They proposed that the breakdown of the resonance of the dipoles in the aromatic amino acids upon treatment with anaesthesia is a major factor responsible for consciousness. However, this hypothesis has not been proven in any biological system. Nevertheless, it is still not known how anaesthesia affects different cellular processes and organelles in a hierarchical and synchronized manner. What are the state-of-the-art functions of different organelles in the cellular environment? Do all organelles react to anaesthesia in a similar manner? To what extent are they altered from their original state? How and whether they recovered after the removal of anaesthesia. Which organelle can survive for a longer duration without disturbing its physical morphology inside the cellular system? Studying the impact of anaesthesia on plants may provide a distinct opportunity to examine the above questions and help to resolve fundamental mechanisms that underlie consciousness.

Here, we studied various organellar dynamics at the apex of living plant roots and the stem connecting to the root upon treatment with both local and general anaesthesia^12,13^. We investigated the dynamics of the mitochondria, endocytic vesicles, chloroplasts, microtubules, and nuclei, which are the main functional organelles in plant cells. The primary goal was to assess the role of ATP, the energy currency of the cell, in the organelles in connection with anaesthesia-dependent conscious awareness. Our results demonstrated that the majority of organelles were highly affected or even damaged by anaesthesia treatment. Interestingly, those mitochondria which permanently damaged are removed from the cell, while those which got depolarize or partially damaged had their membrane potential restored after removal of anaesthesia. some of them regenerated and retained their structure and function after the removal of anaesthesia. In contrast, the nuclei achieved a synchronized arrangement upon treatment with anaesthesia without damaging their overall structure. The synchronized response exemplifies a non-local phenomenon where all nuclei respond identically to external stimuli and carry nucleus-to-nucleus communication in an orchestrated way. The process was completely independent of ATP. A detailed understanding of the chromatin level investigation suggested that euchromatin, the active component of chromatin structure inside the nucleus, reorganized in the nuclear periphery for signal communication from one nucleus to another. The results demonstrated that, out of several organelles, the nucleus survived for a longer duration without disturbing its physical entity. It also serves as the main powerhouse for the cell to survive and helps other organelles to be regenerated. Our findings further demonstrate that chromatin organization and nucleus-to-nucleus communication may serve as potent biomarkers in carriers of conscious awareness.

## Results

We selected two plants (1) tomato (*Solanum lycopersicum*) and (2) brinjal (*Solanum melongena*) for our study because they are easily grown and ideal for study owing to their well-documented physiological information. Tomato and brinjal seedlings were grown under controlled conditions using a commercial plant growth chamber **(Figure S1)**. Local anaesthesia lidocaine, general anaesthesia etomidate and neorof (propofol) were used in our study. Following the reported protocol, 1% lidocaine, 0.2% etomidate, and 1 % neorof were added to the solution of the roots of tomato and brinjal and incubated for an hour in our study ^23–26^. The nuclear arrangement was assessed using DAPI staining, whereas chromatin organization was analyzed by antibody staining of histone proteins. Histone markers were used to visualize euchromatin (H3K4Me3) and heterochromatin (H3K9Me3). While mitochondrial function was evaluated using MitoTracker Green (MTG), Tetramethylrhodamine ethyl ester (TMRE) was used to study the mitochondrial membrane potential (MMP). Autophagy was assessed with the use of LysoTracker Red (LTR), which marks acidic organelles such as lysosomes and autophagosomes. Vesicle trafficking dynamics were monitored using FM4-64, a fluorescent dye that integrates into the vesicle membranes. Microtubule dynamics were studied using Tubulin Deep Red, a dye that specifically labels tubulin proteins. Autofluorescence, which typically originates in a few instances in plants, was monitored during measurement. However, chlorophyll disruption was analyzed via autofluorescence, which naturally emits fluorescence under specific conditions. The reversibility of anaesthetic effects was evaluated by removing anaesthesia and reintroducing normal media. This was done after removing the anaesthesia by washing the sample with PBS buffer four times with a one-hour recovery period. We presented both confocal microscopic images and super-resolved images acquired using super-resolution radial fluctuation microscopy (SRRF) technique ^27^. Image analysis was performed using ImageJ and SPSS (Statistical Package for the Social Sciences).

### Effects of Anaesthesia on the mitochondrial dynamics

The detailed mitochondrial dynamics and MMP were studied by double staining of the plant roots with both MTG and TMRE. Under control conditions without anaesthesia, mitochondria exhibited a stable membrane potential and polarization, which was evident from the highly intense TMRE imaging **(Figure 1a (ii) & S2a (ii))**. Column (i) in Figure 1 shows the MTG staining. The merged image in the last column of **Figure 1a (iv)** shows the colocalization between MTG and TMRE. Anaesthesia treatments caused significant depolarization of mitochondrial membrane, as evidenced by decreased TMRE fluorescence and hence the reduction of MMP **(Figures 1b-d (ii) & S2b-d (ii))**. Depolarized mitochondria are expected to repolarize, but damaged mitochondria must be removed from the cell through autophagy ^28^. Hence, we examined mitochondrial recovery in the next stage. It was evident that depolarized mitochondria became repolarized again **(Figures 1e (ii) & S2e (ii))** after removing the anaesthesia ^29^. Next, we verified whether the mitochondria were damaged upon anaesthetic treatment. This was accomplished by performing an autophagy experiment. To investigate the autophagic response under different anaesthetic conditions, LTR was used to monitor autophagosome production. Autophagy was not detected under the control conditions (**Figure S3a (ii) & S4a (ii)**). However, following anaesthetic treatment, a significant increase in the fluorescence intensity of LTR was observed.

**Figure 1:**
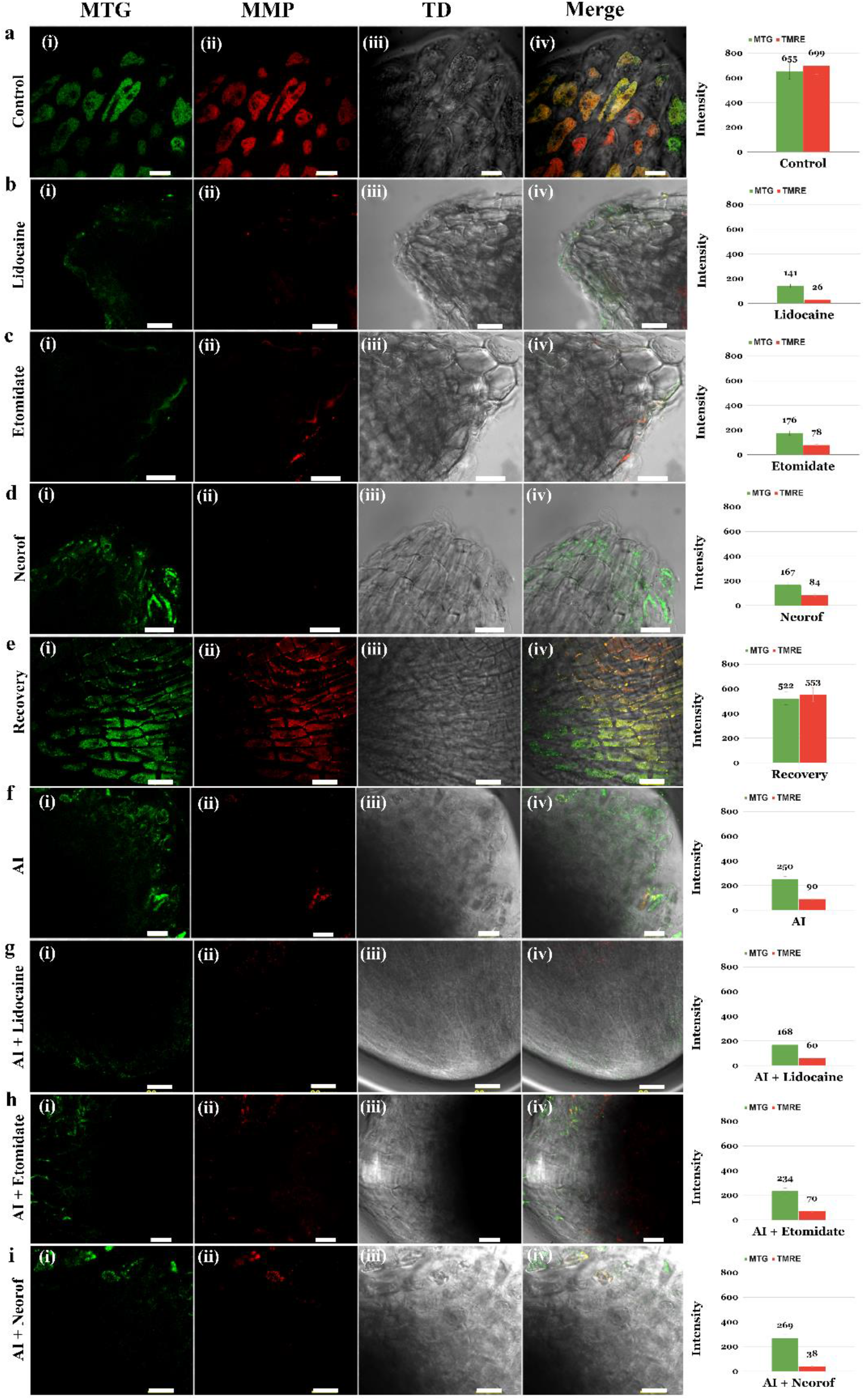
(Tomato) Mitochondrial Dynamics and MMP Under Various Anaesthesia Treatment. The figure displays the effects of different anaesthesia on MMP, **(a)** Control group (without anaesthesia) showing well-polarized mitochondria with high fluorescence intensities of (i) MTG and (ii) TMRE. **(b-d)** represents the mitochondria after various anaesthetic (Lidocaine, Etomidate and Neorof) treatments. The data shows progressively decreased MTG and TMRE fluorescence, thus indicating depolarization or damaged mitochondria. **(e)** The recovery of mitochondria and MMP after removal of anaesthesia **(f)** Treatment with 2 mM sodium azide (ATP inhibitor; AI), resulting in reduced MTG and TMRE fluorescence. This data suggests significant mitochondrial depolarization. **(g-i)** Combined treatment of AI and different anaesthetics. (iii) Transmitted light differential interference (TD) image and (iv) is the merged image of (i), (ii) and (iii). The right-hand side bar graphs are the calculated fluorescence intensity of the images represented in (i) and (ii). The scale bar is 20 µm.

This confirmed the production of damaged mitochondria, along with a corresponding increase in autophagosome formation **(Figures S3b-d (ii) & S4b-d (ii)**). The anaesthesia recovery data suggested the disappearance of the autophagosome, as evidenced by the quenching of the fluorescence intensity of LTR (**Figures S3e (ii) & S4e (ii)**).

Mitochondria are known to be the powerhouses of the cell responsible for ATP production, which leads to various cellular processes. As a result, it could be obvious to understand whether ATP plays any role in mitochondrial function upon treatment with anaesthesia. Sodium azide, a known ATP inhibitor (AI), inhibits cytochrome C oxidase within the mitochondria, thereby reducing ATP generation and disrupting ATP-dependent cellular activity ^30^. **Figure 1f (ii) and S5a (ii)** show a substantial decrease in TMRE intensity, suggesting extensive mitochondrial depolarization. The anaesthetic treatment along with AI led to a more pronounced reduction in MMP **(Figures 1g-I (ii) & S5b-d (ii))**. Interestingly, on the other hand, the combined effect of AI and anaesthesia in comparison to ATP alone showed a substantial increase in autophagy **(Figure S6 (ii) & Figure S7 (ii))**. These findings underscore the role of anaesthesia in enhancing autophagic processes aimed at eliminating dysfunctional mitochondria, as evidenced by the observed data. This observation highlights the critical interplay between mitochondrial dynamics, ATP synthesis, and autophagy in cellular responses to anaesthetic stress. This further emphasizes the significant impact of these treatments on mitochondrial health and cellular homeostasis.

### Anaesthetic effects on the movement of endocytic vesicles in roots

Vesicle trafficking is an extremely important process for cellular communication, because it delivers signaling molecules, nutrients, and minerals between different cellular compartments and the surrounding environment. Under normal conditions, the movement of vesicles within the cells of plant root tips occurs frequently and in significant numbers (**Figure 2a (iii) & Figure S8a(iii)**). However, anaesthetic treatments drastically diminished or stopped the mobility of vesicles, which suggests that there are disruptions in the pathways responsible for cellular signalling (**Figure 2b-d (iii) & Figure S8b-d (iii)**). (Write about Figure 2f also) We further investigated whether inhibition of ATP affected vesicle movement. We found that AI alone had a lesser impact on vesicle trafficking compared to anaesthetics (**Figure 2e (iii) & Figure S8e (iii)**). However, the combination of AI and anaesthetics significantly decreased vesicle movement **(Figures 3a-d (iii) & S8f-h (iii))**. This finding underscores the crucial role of vesicle trafficking in maintaining cellular homeostasis and communication under normal conditions, with disruptions leading to impaired signaling and nutrient transfer ^31^.

**Figure 2:**
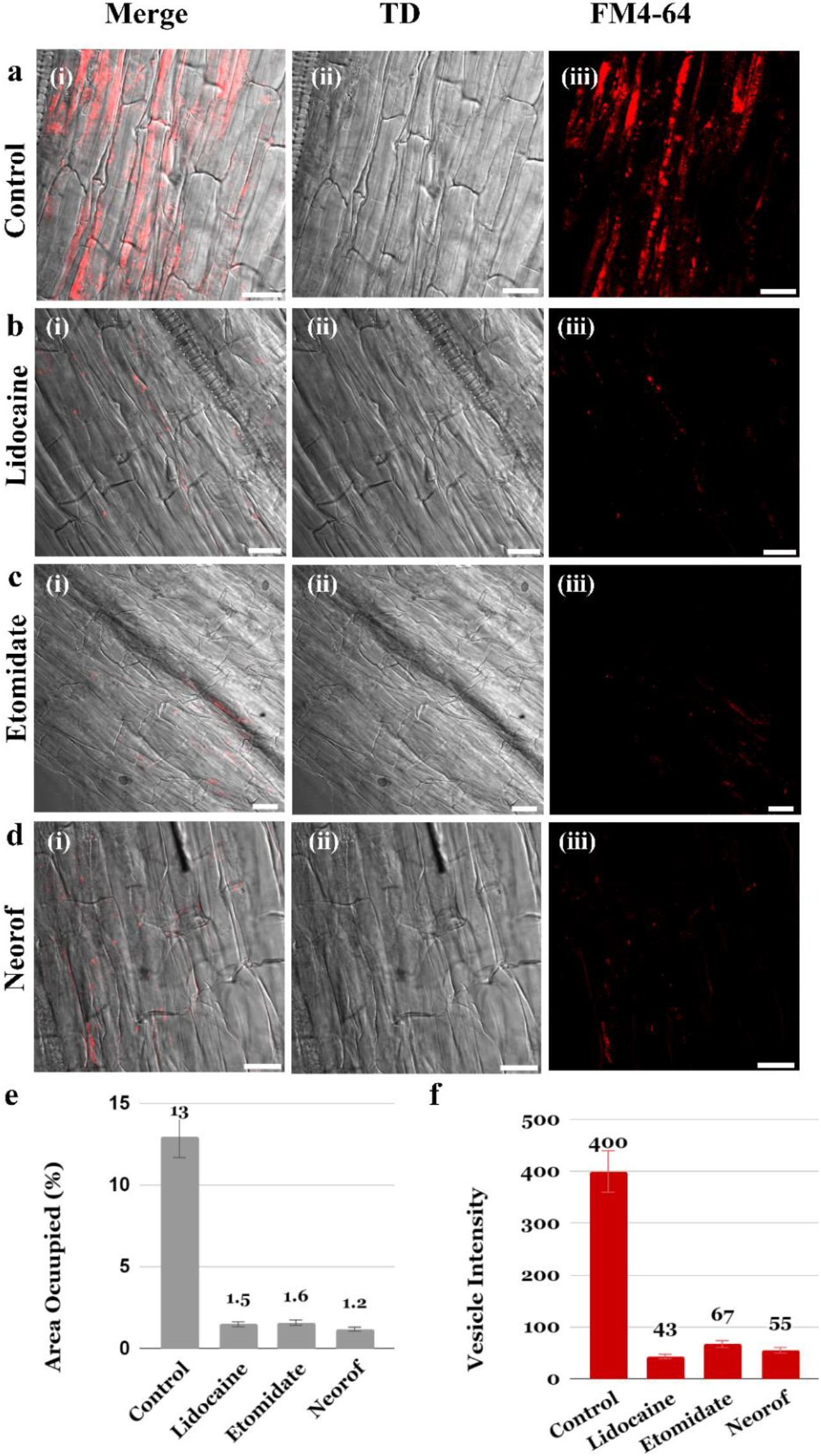
(Tomato) Effects of Different Anaesthetic Treatments on Endocytic Vesicle Trafficking. This figure shows the impact of various anaesthetic treatments on endocytic vesicle trafficking in plant root cells. **(a)** Control condition (without anaesthesia treatment) display (iii) a large fluorescence intensity for vesicle marker FM4-64. This data suggests a large number of vesicles and their robust trafficking. **(b-d)** Reduction in extensive fluorescence intensity (iii) for vesicle marker FM4-64 after treatment with different anaesthetics, suggesting the inhibition of vesicle and their trafficking. (ii) TD image and (i) is the merged image of (ii) and (iii). (**e)** The calculated quantitative area of vesicle trafficking and (f) calculated fluorescence intensity for each image presented in (iii) of (a-d). The scale bar is 20 µm.

**Figure 3:**
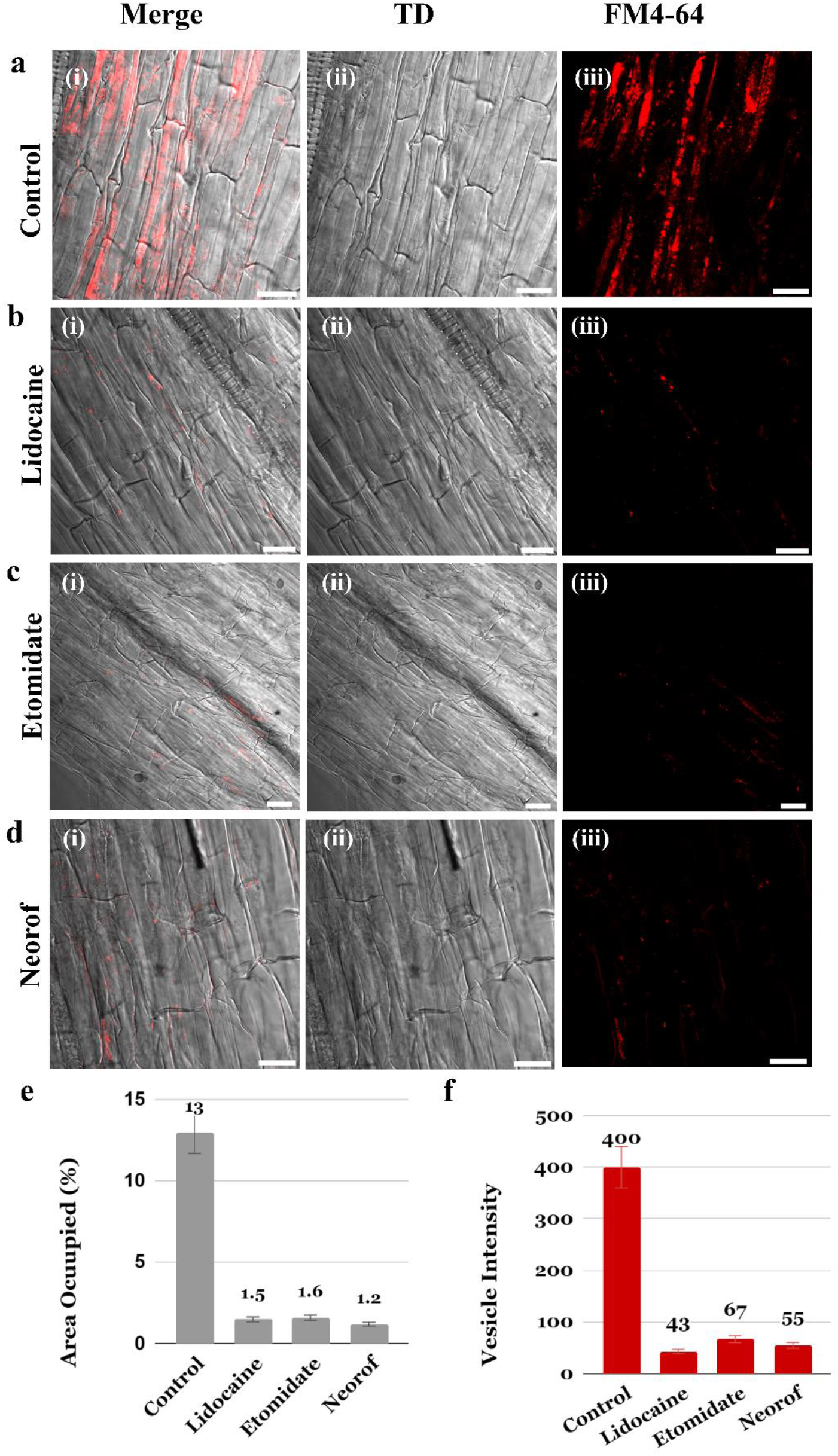
(Tomato) Effects of AI and Combined Effect with Anaesthesia Treatments on Endocytic Vesicle Trafficking. The figure shows the impact of various treatments on endocytic vesicle trafficking in plant cells, with the area occupied by vesicles expressed as mean ± SD for each group and it also shows intensity graph **(a)** Vesicle numbers with AI alone (without anaesthesia), indicating no significant effect of ATP on vesicle trafficking. **(b-d)** A substantial reduction in vesicles numbers and their trafficking was observed upon treatment with anaesthesia. This data suggests a synergistic inhibitory effect. (ii) TD image and (i) is the merged image of (ii) and (iii). (**e)** The calculated quantitative area of vesicle trafficking and (f) calculated fluorescence intensity for each image presented in (iii) of (a-d). The scale bar is 20 µm.

Under control conditions, the average area of the endocytic vesicle movement was approximately 13 % of the total area **(Figure 2e)**. The application of lidocaine resulted in a considerable decrease in vesicle trafficking to a mean value of 1.4 ± 0.8 (p < 0.0001). Both etomidate and neorof caused a considerable decrease in vesicle trafficking. Etomidate reduced it to 1.6 ± 1.4 (p < 0.0001), whereas neorof reduced it to 1.2 ± 0.4 (p < 0.0001). The administration of an AI resulted in a moderate decrease in vesicle trafficking, with an average of 10.7 ± 1.3 (p = 0.06) (**Figure 3e**). When AI was combined with lidocaine, vesicle trafficking was considerably reduced, with a mean value of 1.5 ± 0.7 (p < 0.0001). Similarly, when an AI was paired with etomidate, it yielded an average of 1.4 ± 1.1(p < 0.0001), and when combined with neorof, it produced an average of 1.3 ± 1.5 (p < 0.0001). The information shows that both anaesthesia treatment and combined treatments with AI effectively reduced endocytic vesicle movement compared to control conditions and AI conditions. The fluorescence intensity bar graphs (**Figure 3f**) support vesicle trafficking data. Similar results were obtained in the roots of the brinjal plants **(Figure S8e)**.

### Anaesthesia Alters Microtubule dynamics extensively in plant roots

Our subsequent studies focused on microtubules, which have been suggested to serve as biomarkers of consciousness. Microtubules are crucial components of the cellular cytoskeleton, playing key roles in maintaining cell shape, facilitating intracellular transport, and enabling cell division^32^. Given the ATP-dependent nature of microtubule dynamics, we aimed to investigate whether microtubule disassembly occurs exclusively under anaesthesia or if other environmental conditions could also affect their assembly. Under controlled conditions, we observed normal microtubule formation, appearing as thread-like structures **(Figure 4a & S9a)**. However, treatment with different anaesthetics interfered with microtubule formation, leading to the disassembly of these filamentous structures **(Figure 4b-d & S9b-d)**. The quantitative decrease in the fluorescence image intensity, as presented on the right-hand side of each image, suggesting that anaesthesia effects structural morphology of microtubules. Remarkably, upon removal of the anaesthetics, microtubule formation was reinitiated, suggesting that the effects of anaesthesia on microtubules might be reversible, allowing cells to regenerate their cytoskeletal filamentous structures **(Figure 4e & S9e)**. Furthermore, when sodium azide was used to inhibit ATP production, the microtubules were disassembled. These data suggest that microtubule dynamics are highly dependent on ATP, which disturbs the structure and functions of microtubules, as observed in **Figure S10 & S11**^33^.

**Figure 4:**
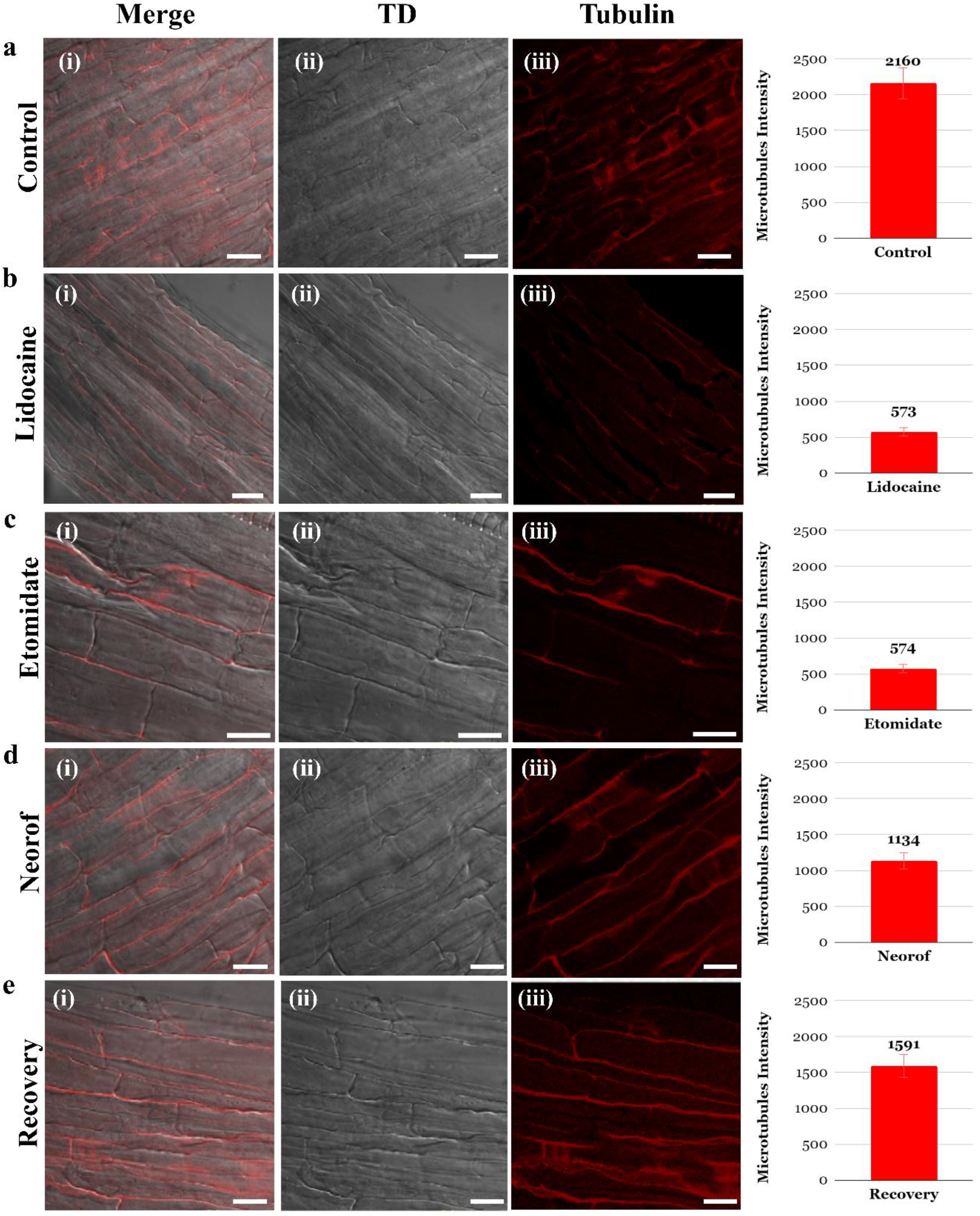
(Tomato) Impact of Anaesthesia on Microtubule dynamics. **(a)** In absence of anaesthesia healthy and thread-like structures is visible as indicated by the high fluorescence intensity of tubulin marker in (iii) **(b-d)** Exposure to anaesthetics results in disrupted microtubule formation, characterized by reduced fluorescence intensity in (iii). **(e)** Upon removal of anaesthesia, microtubule formation resumes, indicating reversibility and an increase in fluorescence intensity. (ii) TD image and is the merged image of (ii) and (iii). The right-hand side bar graphs are the calculated fluorescence intensity of the images represented in (iii). The scale baris 20 µm.

### Impact of Anaesthetics on Chloroplast–Nucleus Association

Communication between the chloroplasts and the nucleus in response to environmental cues is mediated by different signals. Plastid–nuclear complexes make it easier for organelles to communicate with each other directly. This improves the precision of the signals and prevents interference from other sources ^34^. In our study, we observed that under normal conditions, communication between the nucleus and chloroplasts remained intact, as observed in the merged image presented in **Figure 5a (iv) and Figure S12a (iv)**. However, anaesthesia disrupts this communication, likely due to its impact on signal transduction, leading to chloroplast damage **(Figure 5b-d (iv) & S12b-d (iv))**. AI alone does not affect this connection, as some processes in the nucleus and chloroplast do not require ATP **(Figure 5e (iv) & S12e (iv))**. However, the combination of anaesthetics and AI disturbs the connection again, indicating that anaesthesia is the primary disruptor **(Figure 5f-h (iv) & S12f-h (iv))**.

**Figure 5:**
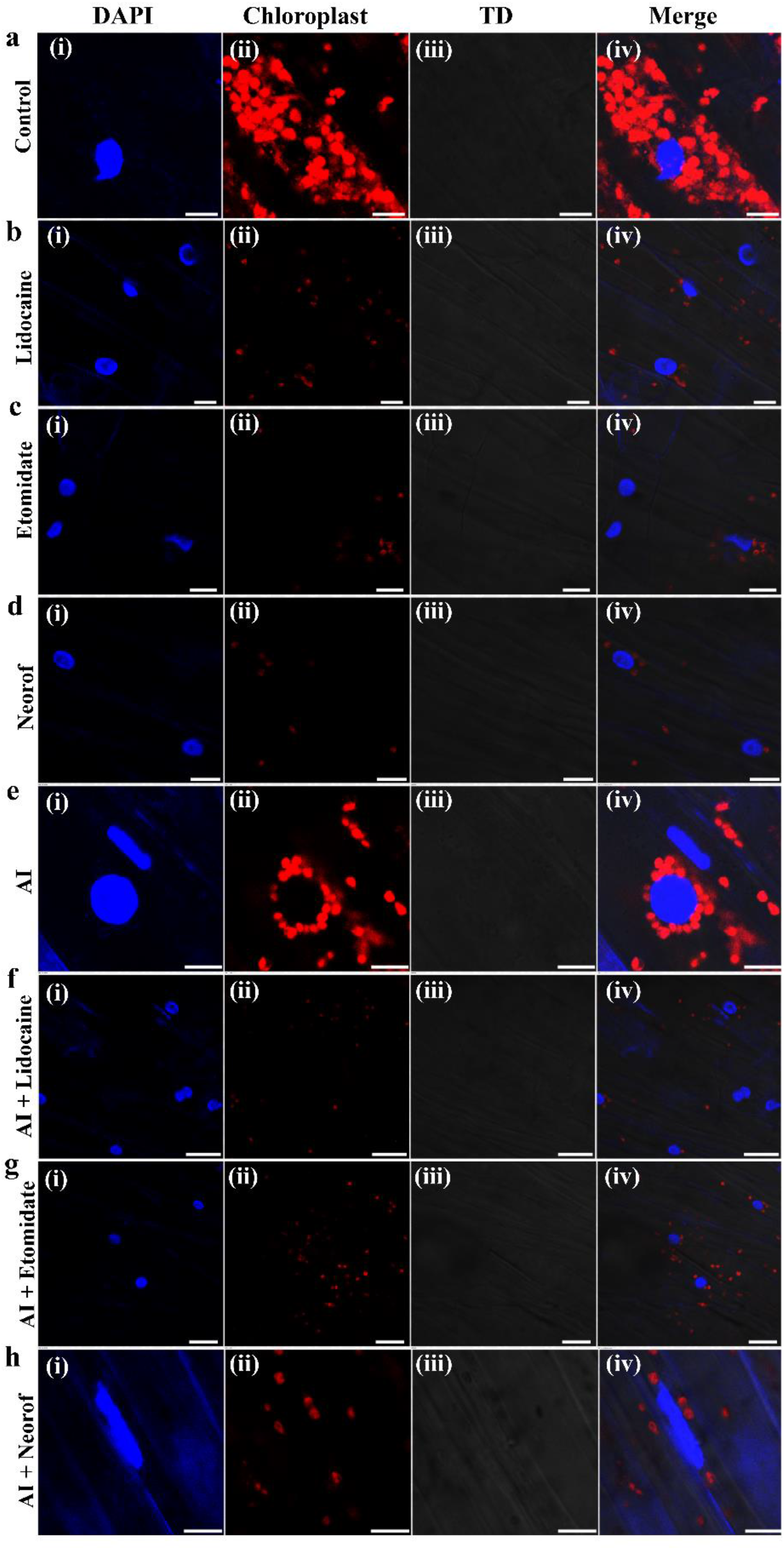
(Tomato) Effects of Anaesthesia and AI on Chloroplast-Nucleus Dynamics in Plant Stem Cells: **(a)** Under normal conditions, a clear association between the nucleus and chloroplasts is observed (iv; merged imageof (i) DAPI in (i)) and (ii) chloroplast. **(b-d)** Anaesthesia disrupts this communication affecting chloroplast integrity as observed in (iv) of (a-d). **(e)** ATP inhibition alone does not affect nucleus-chloroplast communication. **(f-h)** Combined treatment with anaesthesia and ATP inhibition disrupts the normal association between the nucleus and chloroplasts, as evidenced by the altered spatial relationship between these organelles. The scale bar is 10 µm.

Additionally, anaesthesia induces autophagy in plant stem cells, targeting impaired chloroplasts. Under controlled conditions, autophagy was absent, as observed in the merged image of (**Figure 6a (iv))**. However, it significantly increased with various anaesthetics due to disturbed signal transduction, leading to damaged chloroplasts and their association with the nucleus (**Figure 6b-d (iv))**. Interestingly, the anaesthesia recovery data showed no autophagosome formation (**Figure 6e (iv))**. The AI alone did not communicate between the nucleus and the chloroplast (**Figure S13a (iv))**. However, the combined effect of AI and anaesthetics disturbs the signal, causing chloroplast damage and triggering autophagy **(Figure S13b-d (iv))**

**Figure 6:**
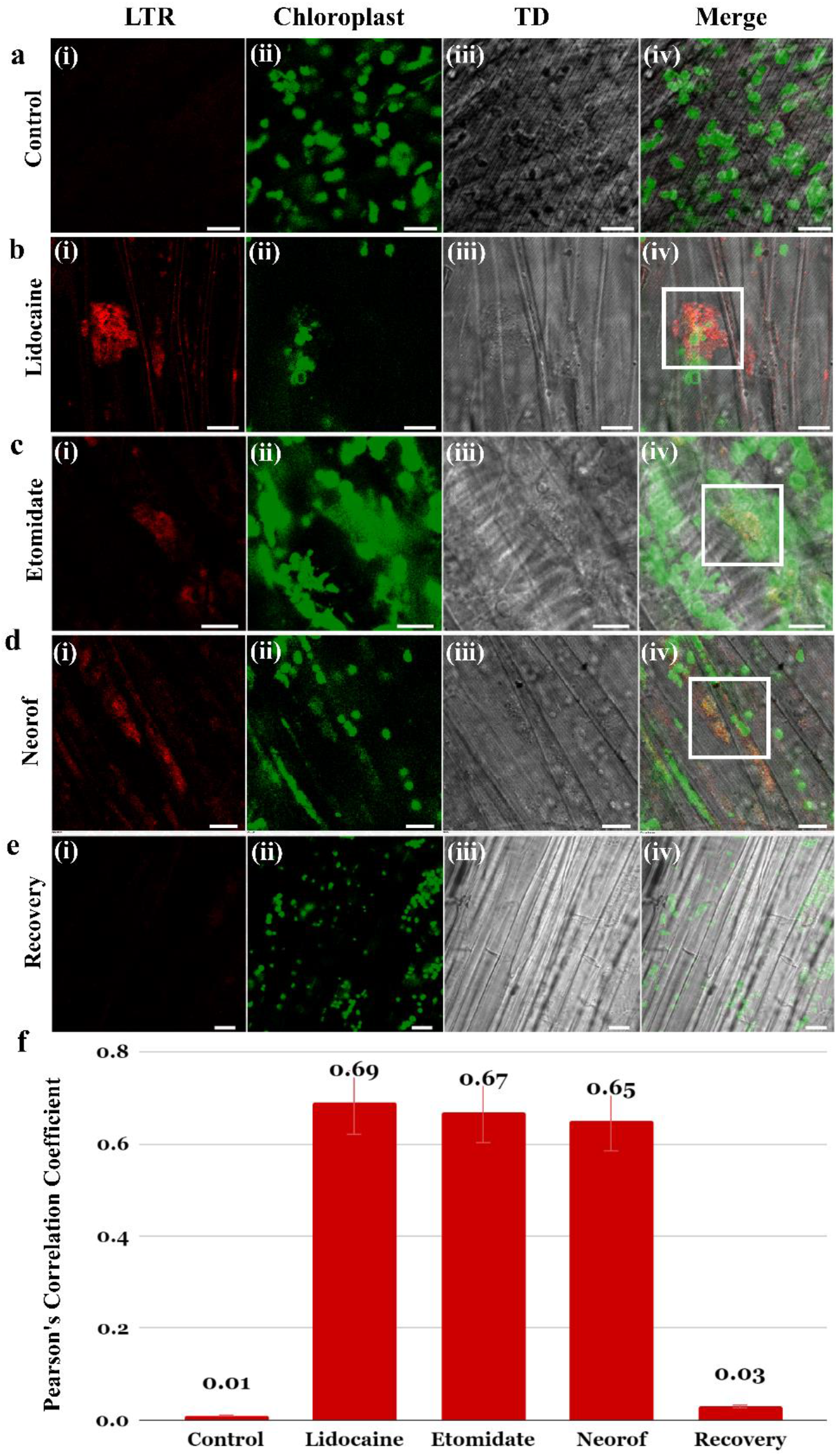
(Tomato) Anaesthesia-Induced Autophagy of Damaged Chloroplasts in Plant Stem Cells: **(a)** Under control condition (without anaesthesia), no autophagosome formation is observed as evident by the negligible fluorescence intensity in LTR channel (i). The merged image of (i) LTR and (ii) chloroplast in (iv) further showed no correlation in between chloroplast and autophagosome. **(b-d)** Different anaesthesia treatments induce varying extents of autophagosome formation around damaged chloroplasts as evident by the autophagosome formation stained by LTR (i) and the merged image of it with the chloroplast in (iv). **(e)** Recovery of chloroplast integrity is observed after the removal of anaesthesia. **(f)** Pearson’s coefficient is used to quantify colocalization, with mean values (bars) and error bars further supports our claim as presented in (a-e). The scale bar is 10 µm.

To examine chloroplast dynamics, we performed an experiment on plant cotyledons. Plastids such as chloroplasts are essential for energy generation in plants, with chloroplasts specifically responsible for energy production. Different Plastid types can undergo conversion in response to changes in metabolism or the environment. Photomorphogenesis is activated by light, while skotomorphogenesis is activated by the absence of light. Chloroplasts originate from proplastids located in the meristems. However, in the absence of light, they transform into etioplasts, which exhibit distinct differences in terms of inner membrane organization and pigment content ^35^. The data presented in **Figure S14a & S15a** shows normal and healthy chloroplasts without anaesthesia treatment. This suggests that anaesthesia affects the perception of light and causes chloroplasts to revert to an etioplast state. Surprisingly, the ATP inhibition results showed healthy chloroplasts, suggesting that ATP inhibition has no negative impact on the chloroplast structure **(Figure S14e, S15e)**. This is likely because chloroplasts carry out photosynthesis and contribute to the production of ATP in the plant system through respiration. Therefore, inhibiting ATP does not substantially affect chloroplasts. When anaesthetics and ATP inhibition were used together, again etioplast-like structures were formed **(Figure S14f-h & S15f-h)**. The combined impacts increased the negative impact on the integrity and functionality of the chloroplasts. The results showed that while anaesthesia disrupts chloroplast integrity, ATP inhibition alone does not affect chloroplast morphology.

### Anaesthetic Effect on Nuclear Arrangement in Plant Root Cells

The nucleus provides valuable information regarding cellular organization, gene regulation, developmental processes, and stress responses. This current observation can help in understanding the fundamental mechanisms of cellular function and how they are influenced by various factors. In our experiment, the nuclei of plant root cells were randomly distributed throughout the cytoplasm under control conditions **(Figure 7a & S16 (I) a)**. After an hour of anaesthetic treatment, the nuclei were rearranged into clear patterns in their original place. These findings suggest that anaesthetics induce the organization of nucleus in a certain manner **(Figures 7b-d & S16 (I) b-d)**. Exposure to 2 mM AI led to random distribution of the nucleus, indicating that ATP does not affect the arrangement of the nuclei **(Figure 7e & S16 (I) e)**. Surprisingly, the combined treatment with AI and different anaesthetics successfully restored the structured nuclear patterns **(Figures 7f-h & S16 (I) f-h)**, suggesting that anaesthetics has the ability to promote nuclear organization. Zoomed image of the nucleus arrangement in synchronized manner under different anaesthetics shown in **Figure S17 a-f**. After removal of anaesthesia, time- dependent recovery of nucleus also observed (**Figure S18 (i)- (iv)**).

**Figure 7:**
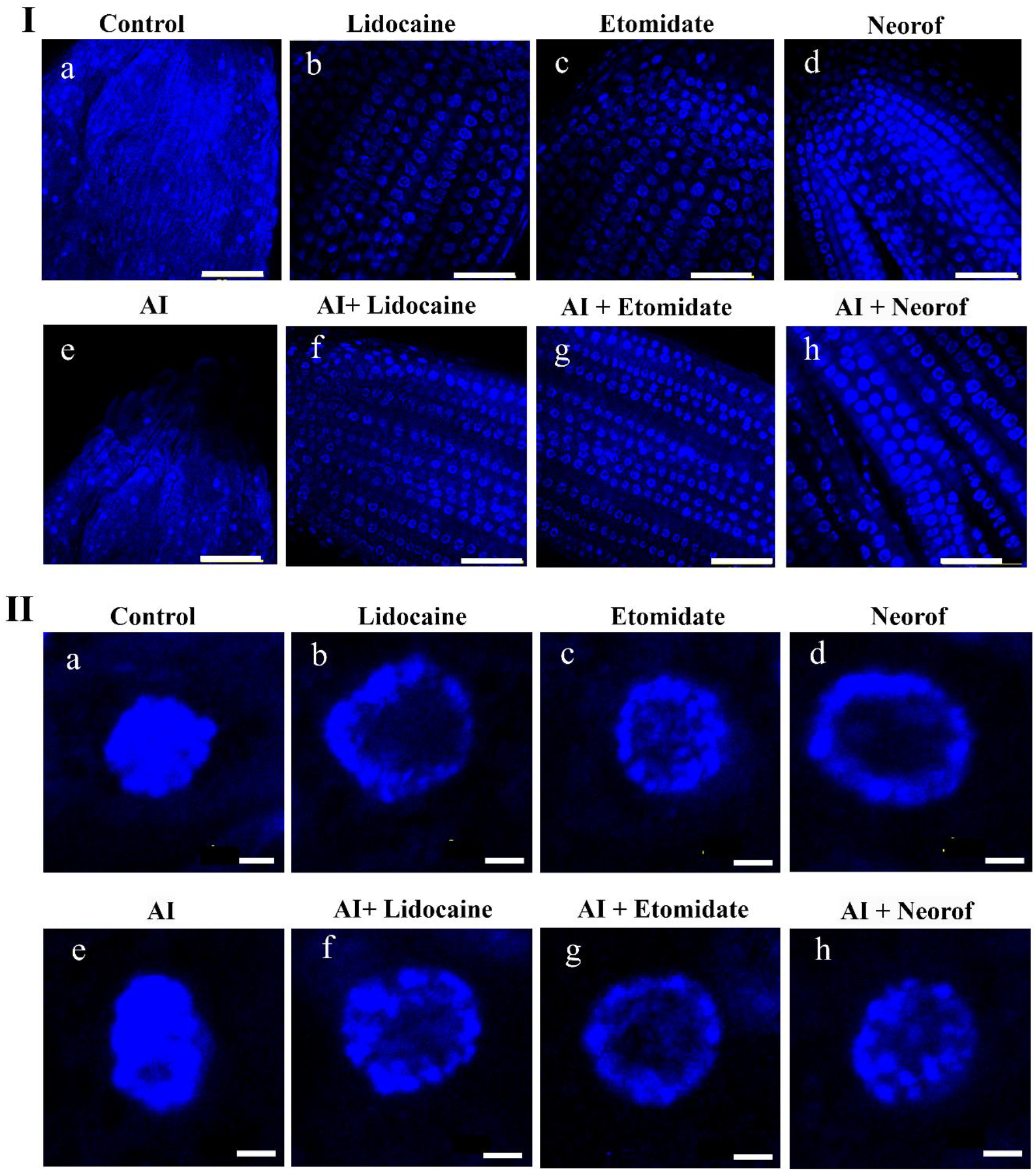
(Tomato) Impact of Different Treatments on Nuclear Arrangement in Plant Root Cells. This figure highlights the effects of various treatments on the arrangement of nuclei in plant root cells. **I (a)** Displays scattered nuclei under normal conditions, indicating a typical, chaotic distribution. **(b-d)** Shows nuclei arranged in specific patterns following treatment with the anaesthetics, suggesting that these compounds induce a reorganization of nuclear positioning. **(e)** AI condition (without anaesthesia) depicted the scattered nuclei similar to the control condition, indicating that ATP ingibitor alone does not significantly alter nuclear arrangement. **(f-h)** Demonstrates nuclei arranged in distinct patterns even in presence of ATP inhibior. (Scale bar represents 10 µm). **II**. Shows zoomed images of a single nucleus of (I) after anaesthetic treatment, most probably highlighting changes in chromatin arrangement. These images provide a closer look at how anaesthetics affect the internal organization of nuclear material. (Scale bar represents 2 µm)

Within the nucleus, chromatin plays a major role in cellular functions. Chromatin is tightly coiled DNA and histones proteins that constitute the majority of the nucleosome. A nucleosome is 147 bp of DNA wrapped in 1.67 left-handed twists around an H2A, H2B, H3, and H4 histone octamer ^36^. Heterochromatin, a compact, transcriptionally inactive structure of chromatin found in pericentric regions, telomeres, and other suppressed areas. On the other hand, in gene-rich genomic regions, euchromatin is more open and transcriptionally active. Chromatin remodeling that alters gene expression allows plants to survive in adverse environments ^37^. When extremely condensed, the chromatin architecture prevents transcription factors, polymerases, and other nuclear proteins from reaching DNA. Stress signals unlock DNA by altering chromatin structure. Chromatin remodeling shifts, reorganizes, repositions, or modifies histones ^38^.

To better understand euchromatin and heterochromatin dynamics, we performed super-resolution radial fluctuation (SRRF) microscopy. This is a powerful tool for investigating cellular dynamics down to the nanometer level by breaking the diffraction limit of light. (cite SRRF article here) The nuclear region typically displays a dispersed and diffuse euchromatin distribution pattern under control condition **(Figure 8I, a)**. Because euchromatin is normally linked to gene-rich regions of the genome that are accessible for transcription, this distribution implies an active transcriptional state ^8,39^. Different anaesthetics cause extensive rearrangement of euchromatin. Euchromatin is mainly localized to the periphery of the nucleus after anaesthetic treatment. **Figure 8I, b-d**, shows that changes in the chromatin state or transcriptional activity may have caused this nuclear architectural alteration ^8^. To determine whether ATP is necessary for euchromatin reorganization, cells were treated with AI. In the absence of ATP, euchromatin remained scattered throughout the nuclear region, similar to the distribution observed under control conditions **(Figure 8I, e)**. When cells were subjected to both ATP inhibition and different anaesthetic treatment, euchromatin once again exhibited a peripheral arrangement within the nucleus **(Figure 8I, f-h)**. Interestingly, regardless of the treatment, heterochromatin remained unaffected. Recent findings suggest that under stress conditions, heterochromatin, typically bound to peripheral Lamina-Associated Domains (LADs) and associated with gene silencing, can dissociate from these regions. This dissociation allows euchromatin, which contains active genes, to occupy these sites, potentially enabling the activation of stress-responsive genes. This dynamic chromatin repositioning highlights the role of cellular adaptation and responses to environmental stressors ^40^.

**Figure 8:**
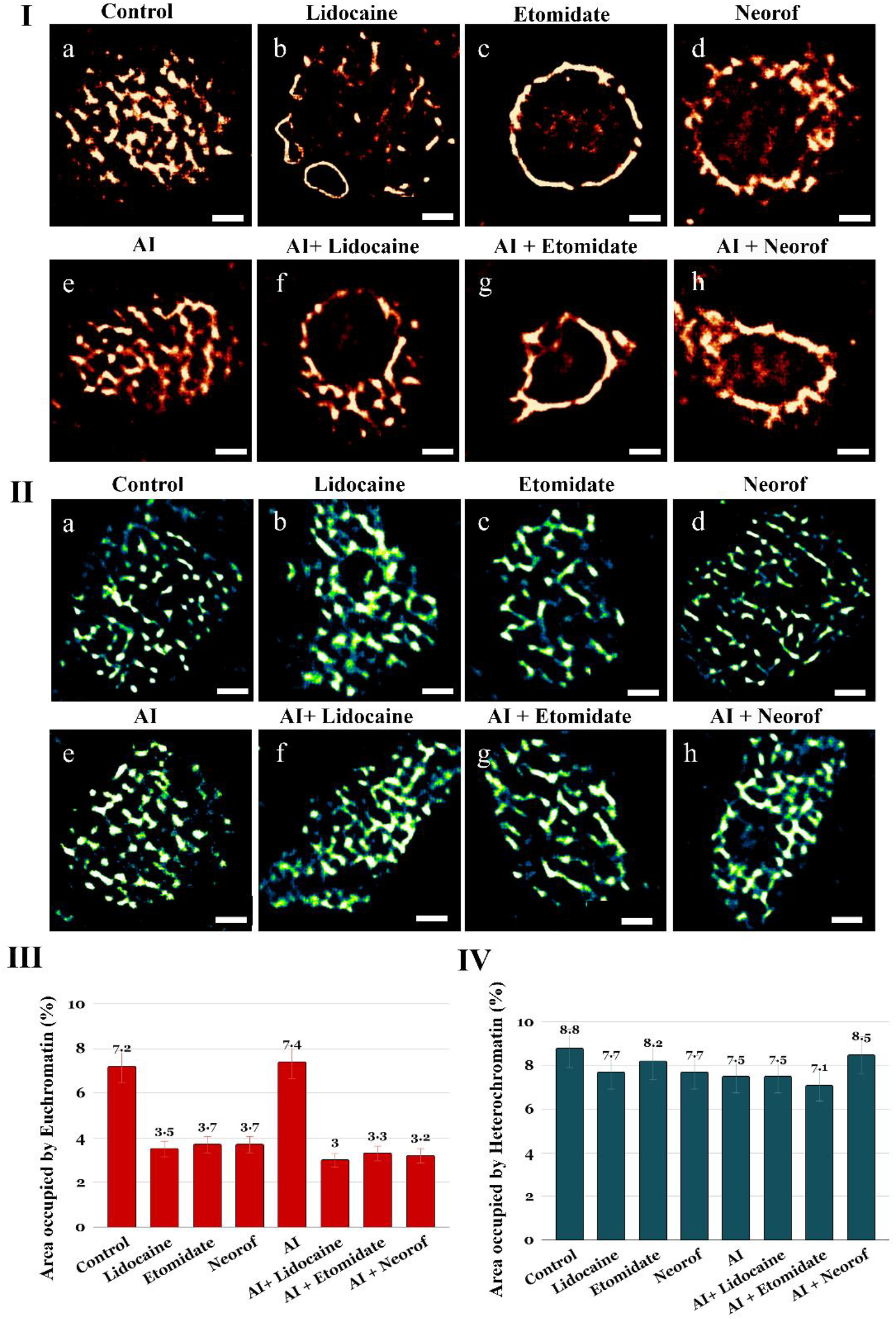
(Tomato) Effect of Different Treatments on Euchromatin and Heterochromatin Distribution in Plant Root Cells. The figure explores how various treatments impact the distribution of euchromatin and heterochromatin in plant root cells. **I (a)** Shows scattered euchromatin, indicating a typical, non-specific distribution within the nucleus. **(b-d)** Displays euchromatin arranged at the nuclear periphery following treatment with these anaesthetics, suggesting that these compounds influence euchromatin positioning. **(e)** Illustrates scattered euchromatin like the control condition (without anaesthesia) but upon ATP inhibition. This result indicates that AI alone does not significantly alter euchromatin distribution. **(f-h)** Demonstrates euchromatin arranged at the nuclear periphery, similar to panels B-D, indicating that the combined treatment also impacts euchromatin positioning. **II**. Shows that heterochromatin distribution remains unaffected across all treatments. This suggests that these treatments specifically influence euchromatin without altering heterochromatin organization. **III & IV**. Shows the area occupied by euchromatin and heterochromatin, expressed as mean (bar) ± SD (error bar) for each group. This quantification provides a comparative analysis of chromatin distribution under different treatment conditions. (Scale bar for SRRF image is 2 µm).

## Discussion

We used confocal microscopy and super-resolution microscopy imaging techniques to investigate plant cell responses to anaesthesia. This study provides new insights and direct evidence for plant consciousness and awareness. Our study demonstrated that anaesthesia significantly affects various cellular processes and structures in plant cells. Anaesthesia depolarizes mitochondria, reduces MMP, and damages several mitochondria under the given experimental conditions **(Figure 1a (ii) & S2a (ii))** ^41^. Autophagy degrades defective mitochondria and restores cellular homeostasis after a disturbance **(Figures S3b-d (ii) & S4b-d (ii)**) ^42^. AI reduces mitochondrial activity and membrane capacity by inhibiting ATP generation. Once AI has been applied to plant cells, anaesthesia has very little role in mitochondrial dynamics, as was observed in the non-ATP-dependent process (**Figure 1f (ii) and S5a (ii))**. After the combined effect of ATP and Anaesthesia, the results showed that membrane potential was completely diminished **(Figures 1g-I (ii) & S5b-d (ii))**.

On the other hand, vesicle movement, essential for cellular communication and nutrient transport, is significantly reduced by anaesthetic treatment, disrupting cellular signaling pathways (**Figure 2b-d (iii) & Figure S8b-d (iii)**)^14^. AI alone had a lesser impact on vesicle trafficking (**Figure 2e (iii) & Figure S7e (iii)**), but their combination with anaesthetics markedly decreased vesicle movement **(Figures 3a-d (iii) & S8f-h (iii))**. Certain cellular processes have backup mechanisms or alternative energy sources that partially compensate for the loss of ATP, allowing for a degree of vesicle movement even under ATP inhibition, as observed in our study (**Figure 2b-d (iii) & Figure S8b-d (iii)**). It has been proposed that anaesthesia disrupts cell membranes and ion channels, leading to altered ion fluxes and membrane potentials. This disruption impairs the function of the cytoskeleton and motor proteins, which are essential for vesicle transport ^43^. The loss of signal transduction and nutrient transfer suggests disruptions in cellular communication pathways that may alter plant consciousness and responses to environmental stimuli ^14^. This finding supports the idea that awareness requires strong cellular signaling.

Microtubules, which are essential for maintaining cell shape and facilitating intracellular transport, exhibit ATP-dependent assembly and disassembly. Most plants and animals have similar microtubule structures. Anaesthetics disrupted microtubule formation, leading to a decrease in fluorescence intensity and thread-like structure as seen in control condition **(Figure 4b-d & S9b-d)**, which reverses upon anaesthesia removal **(Figure 4e & S9e)**. AI similarly causes microtubule depolarization, and their combined use with anaesthetics underscores the crucial role of ATP in microtubule dynamics **(Figure S10 & S11)**. Our results indicate that anaesthesia significantly disrupts microtubule formation and promotes low fluorescence intensity, suggesting that microtubules may not solely mediate consciousness. This finding highlights the complexity of microtubule stability and the broad influence of cellular mechanisms. Further research is needed to explore the detailed cellular responses to stress and their implications for consciousness ^32 33^.

Our study highlights the impact of anaesthesia on nucleus-chloroplast communication and autophagy in plant stem cells. Under normal conditions, this communication was intact, as shown in **Figures 5a(iv) and S12a(iv)**. Anaesthesia disrupts this connection, likely through impaired signal transduction, leading to chloroplast damage **(Figures 5b-d(iv) & S12b-d(iv))**. AI alone does not affect this communication, but when combined with anaesthesia, they do, indicating anaesthesia as the primary disruptor **(Figures 5f-h(iv) & S12f-h(iv))**. We also observed that anaesthesia-induced autophagy by targeting damaged chloroplasts. Autophagy was absent under control conditions **(Figure 6a(iv))** but increased significantly with anaesthesia due to disrupted signal transduction **(Figures 6b-d(iv))**. Upon anaesthesia removal, autophagy ceased, suggesting reversibility **(Figure 6e(iv))**. While ATP inhibition alone does not affect nucleus-chloroplast communication **(Figure S13a(iv))**, combined with anaesthesia, it disrupts signaling, causing chloroplast damage and triggering autophagy **(Figures S13b-d(iv))** ^34,44^. Anaesthesia also causes chloroplasts to become etioplast-like. Anaesthesia induced etioplast-like changes in chloroplasts, affecting their morphology and function (**Figure S14b-d and S15b-d)**. ATP inhibition alone does not affect chloroplast integrity, suggesting a little role in this process. However, combined with anaesthesia, it exacerbates chloroplast damage, emphasizing the primary disruptive role of anaesthesia **(Figure S14f-h & S15f-h)**. ^44,45^. The complex regulation of chloroplast activity is demonstrated by changes in light perception and organelle differentiation.

The most striking discovery was that anaesthesia produced substantial changes in the composition of the plant cell nucleus and chromatin (**Figure 7, 8)**. It aligns with the results obtained in animal research studies showing the repositioning of chromatin towards the periphery of the nucleus ^46^. This suggests changes in transcriptional activity and gene expression patterns **Figure 8I, b-d**. This observation aligns with the idea that the nucleus plays a crucial role in consciousness, as the accumulation of euchromatin near the nuclear envelope might sustain signaling and transcription processes essential for maintaining basic awareness in plants ^8^. Despite the significant role of ATP in cellular metabolism, ATP inhibition did not affect nuclear or chromatin organization, indicating that consciousness may involve mechanisms other than ATP-dependent activity **(Figure 7,8, e-h)**.

Chromatin remodelling activities involve two enzymatic mechanisms: ATP hydrolysis changes DNA-histone interactions, and specialized enzymes methylate DNA or alter histone residues covalently ^47^. Dynamic changes in chromatin composition and organization may cause stress-induced transcriptional alterations; however, plant stress responses are correlated ^8^. Studies have shown that due to stress, chromatin remodelling occurs to maintain cellular activities and proper signalling in cells. For example, bivalent H3K4me3 and H3K27me3 marks, related to active genes in response to cold stress, improve chromatin accessibility, allowing regulatory proteins to access gene expression ^48^. A recent study demonstrated that DMRs and several DEGs in bok choy (Brassica rapa) and rice (Oryza sativa) under heat, drought, and salinity stress indicate how DNA methylation affects gene expression ^49^. Many pharmacological, biophysical, and genetic studies have examined chromatin remodeling and how remodelers regulate cellular functions under stress.

In our study, euchromatin was repositioned under different anaesthetic conditions, but heterochromatin was not affected. Repositioning did not occur when ATP production was inhibited. Unlike most other studies, our study found only basic relocation and no ATP-dependent remodeling during stress. As in animal cells (Drosophila larvae), plants may rearrange chromatin at the nuclear periphery in response to signals ^46^. Recent research has shown that nuclear peripheral heterochromatin stabilizes the genome and silences genes. Heat stress disrupts tomato chromatin and constitutive heterochromatin connections, which may be deleterious; however, it also promotes dynamic chromatin changes that change gene expression to adapt to environmental stress ^40,50^. CRWN1, CRWN4, and KAKU4 proteins in the plant nuclear lamina dynamically respond to stress. Each component moves to the periphery of the nucleus. These components redistribute to the nucleoplasm during heat stress without affecting the nuclear size or shape, suggesting that their structural role in nuclear morphology predates the interphase. Heat stress breaks the nuclear lamina and moves CRWN1, CRWN4, and KAKU4 to the nucleoplasm. CRWN1chromatin interactions and gene expression affect disassembly. After posttranslational modifications, plant nuclear lamins may recognize and respond to stress ^51^. Most cells shut down during anaesthesia and laminal proteins disintegrate, removing heterochromatin from the periphery and repositioning euchromatin. Euchromatin moving to the periphery may affect gene expression and transcription because most of the transcription and RNA processing occurs in the peripheral area ^46^.

Interestingly, although ATP plays a significant role in the metabolic processes of cells, the use of AI does not result in nuclear or chromatin organization in plants. Although the restoration of chromatin organization following anaesthesia with AI reveals that consciousness may entail mechanisms that go beyond ATP-dependent cellular activities, this demonstrates that there are other mechanisms at play in the process of preserving fundamental awareness in plants. It has been suggested that the nucleus may act as a carrier of consciousness owing to the accumulation of euchromatin, which was detected in close proximity near to the nuclear envelope after anaesthesia treatment. Euchromatin, the active chromatin responsible for many cellular functions, is rearranged in the peripheral region to sustain signalling and transcription processes, which may potentially preserve the fundamental awareness that exists within an organism. A conclusion that may be drawn from this is that although anaesthesia does interfere with cellular activity, particularly in the nucleus, it does not totally eliminate cellular functions.

In summary, the effects of anaesthesia on plants revealed that anaesthetics significantly alter plant cellular processes, such as mitochondrial function, autophagy, vesicle trafficking, and microtubule dynamics. Anaesthesia induced substantial changes in chromatin and nuclear arrangement, suggesting a critical role in maintaining fundamental awareness, even under stress. Interestingly, ATP inhibition did not affect the nuclear or chromatin organization, highlighting the nucleus and chromatin as potential carriers of consciousness. Overall, these findings suggest that nuclear and chromatin arrangement are vital for preserving basic cellular functions and consciousness during anaesthesia. Therefore, fundamental consciousness may persist even under anaesthetic conditions because of the resilience of these organelles.

## Conclusion

This study has provided crucial insights into the effects of anaesthesia on plant cellular processes, particularly focusing on the root apex of tomato (*Solanum lycopersicum*) and brinjal (*Solanum melongena*) seedlings. Our data suggest that mitochondria, microtubules, and endocytic vesicles were highly altered or damaged under the same experimental conditions, and their functions also majorly depended on ATP. In contrast, our findings revealed that anaesthesia-induced euchromatin relocation to the nuclear periphery occurs in a non-local, ATP-independent manner, without disturbing much of its structure. Nucleus-to-nucleus communication (internuclear communication) could be clearly visible upon treatment with anaesthesia. This highlights the nucleus and chromatin organization as essential carriers of consciousness, underscoring the sophisticated capability of plant cells to maintain homeostasis and conscious behavior akin to human brain function. We demonstrated that chromatin repositioning and architecture serve as significant biomarkers of plant consciousness. These results contribute to a broader understanding of consciousness and its cellular mechanisms in various biological systems.

## Methodology

### 1. Selected plants and growth conditions

The plant material consisted of tomatoes *(Solanum lycopersicum)* and brinjals (*Solanum melongena)*. After sterilization with 70% ethanol, seeds were carefully rinsed with sterile distilled water. Germination was performed on ½ Murashige and Skoog (MS) agar plates. The plates were sealed to preserve sterility, and the seedlings were grown under controlled conditions at 21°C with 16 h of light and 8 h of darkness. In this experiment we used 2-week-old tomato and brinjal seedlings.

### 2. Treatment Strategies for Anaesthesia

Seedlings were treated with several anaesthesia agents including Lidocaine (1%), Etomidate (0.2%), Neorof (1%), and the ATP inhibitor sodium azide (2 mM) for 1 hour. We took 1 hr anaesthetic treatment because most of the paper suggests that 1 hr anaesthetia treatment is enough to make living being unconscious. Post-treatment, the seedlings were subjected to a recovery process by removing the anaesthesia and washing with PBS buffer three times, followed by a 1-hour rest period. Observations were then made using confocal microscopy to assess recovery.

### 3. Effect of Anaesthesia on Nuclear Arrangement

Root cells nuclei were stained with DAPI (1 mg/mL; Thermo Fisher) and washed with phosphate-buffered saline (PBS). After washing with PBS, root cells were subjected to different treatments. Chromatin arrangement was assessed through immunofluorescence staining. Various anaesthetic treatments were applied to the seedlings, including lidocaine (1%), etomidate (0.2%), neorof (1%), the ATP inhibitor sodium azide (2 mM), and the combined effect of the ATP inhibitor and anaesthesia. The cells were incubated overnight at 4°C with primary antibodies (1:100 H3K4Me3 and H3K9Me3; Abclonal), washed with PBS containing 0.1% Tween 20, and incubated with a secondary antibody (1:600 Cy3-conjugated; Abclonal) for 1 h. After the final washes, coverslips were mounted with glycerol and sealed before imaging.

### 4. Effect of Anaesthesia on Mitochondria and Mitochondrial Membrane Potential

Root cells mitochondria were labelled with MitoTracker Green, a fluorescent dye manufactured by Thermo Fisher Scientific and washed with PBS. Following repeated PBS washes, several anaesthetic treatments were administered to seedling roots. These treatments included lidocaine (1%), etomidate (0.2%), neorof (1%), the ATP inhibitor sodium azide (2 mM), and the combined impact of an ATP inhibitor and anaesthesia. The membrane potential was visualized using tetramethylrhodamine ethyl ester (TMRE; Thermo Fisher).

#### Autophagy Experiment for Mitochondria

Lysosomes were identified by staining with LysoTracker Red (Thermo Fisher Scientific) in root cells. Following the staining process, root cells were subjected to the following treatments: 2% lidocaine, 0.2 percent etomidate, 1 percent neorof, 2 mM sodium azide, an ATP inhibitor, and the combined action of an ATP inhibitor and anaesthesia. Following the remoFval of the anaesthetic and subsequent introduction of normal media, the reversibility of the effects of anaesthesia was evaluated.

### 5. Endocytic Vesicle Trafficking

Tomato seedlings were stained with FM4-64 dye (Thermo Fisher Scientific) and exposed to 2% lidocaine for 30 minutes. To prevent the recycling of endocytic vesicles, further treatment with Brefeldin A (BFA; Thermo Fisher) was administered for a period of thirty minutes. Over the course of the experiment, the seedlings were subjected to a number of different Anaesthesia treatments. These treatments included lidocaine (1%), etomidate (0.2%), neorof (1%), the ATP inhibitor sodium azide (2 mM), and the combined impact of an ATP inhibitor and Anaesthesia.

### 6. Microtubules Intensity

Seedlings were stained with a tubulin deep red marker (Thermo Fisher) and exposed to different treatments for a 6-hour incubation period to visualize microtubules in the root apex. The treatments included lidocaine (1%), etomidate (0.2%), neorof (1%), the ATP inhibitor sodium azide (2 mM), and the combined effect of an ATP inhibitor and Anaesthesia.

### 7. Effect of Anaesthesia on Chloroplast Disruption

Chloroplast dynamics were observed by microscopy before and after exposure to assess the impact on chloroplasts in the tomato seedling leaves. Chloroplast autofluorescence was used for visualization. Various anaesthetic treatments were applied to the seedlings, including lidocaine (1%), etomidate (0.2%), neorof (1%), the ATP inhibitor sodium azide (2 mM), and the combined effect of an ATP inhibitor and anaesthesia.

### 8. Impact of Anaesthesia on Nucleus-Chloroplast Association

The root cells were stained and washed with PBS 3 times. Nuclei were stained with DAPI. The study assessed changes in chloroplast and nucleus arrangement in plant cells under the influence of Various anaesthetic treatments were applied to the seedlings, including lidocaine (1%), etomidate (0.2%), neorof (1%), the ATP inhibitor sodium azide (2 mM), and the combined effect of an ATP inhibitor and anaesthesia.

### 9. Chloroplast Autophagy Experiment

Root cells were stained with LysoTracker Red, and subjected to various treatments. The reversibility of the effects was assessed by removing Anaesthesia and reintroducing the normal media. Various anaesthetic treatments were applied to the seedlings, including lidocaine (1%), etomidate (0.2%), neorof (1%), the ATP inhibitor sodium azide (2 mM), and the combined effect of an ATP inhibitor and anaesthesia.

### 10. Confocal / SRRF Microscopy and Image Processing

#### a. Confocal Microscopy

Nikon Eclipse Ti inverted microscope was used for the confocal microscopy and images were acquired using Nikon Nis-Element software. The cell samples were excited by a 639 nm laser, and the emission was collected using the appropriate filter sets. The colocalization study was performed using a 60x (1.40 NA) oil immersion objective.

#### b. SRRF Bioimaging of Euchromatin and Heterochromatin

Euchromatin and heterochromatin structures were super-resolved using the SRRF algorithm, to visualize and achieve higher-order structures in the Nikon Ti Eclipse microscope with 100X Plan apo λ oil immersion objective (NA 1.45) and then magnified with 1.5x magnification. Andor iXon ultra 897U EMCCD camera was used for image projection. The acquisition procedure was monitored with Andor Solis software having camera setup of 50ms (∼20 fps) exposure time, readout rate 17 MHz at 16-bit, shift speed 3.3, along with gain 3 of the preamplifier. Based on the required shape and sizes frames were cropped. Movies of 5,000 staked images were recorded along with 1000 stacked images in TD mode of the same area, then all the files were saved in .fits file format for additional SRRF processing. To excite the histone mark labeled fluorophores, a fluorescent lamp linked to the microscope was used. Appropriate emission/excitation filter sets were chosen based on the fluorophore of interest. We used Nikon’s perfect focus system (PFS) to take the videos for a longer time to avoid defocusing on the target object area.

#### c. SRRF Image Reconstruction for Chromatin

In order to break the diffraction limit, Andor Solis’s acquired movies (3000-5000 stacked images) of each sample set were analyzed by implementing an open-source version of NanoJ-SRRF on a high-performance NVIDIA GeForce RTX 3070 GPU. Then NanoJ-core plugin of SRRF was used for the drift correction to remove the effect of drift in both the x and y-axis. Furthermore, NanoJ-SRRF plugin of ImageJ was used to reconstruct the super-resolved images by following the method above mentioned. Our optimized parameters viz the ring radius 0.5, radiality magnification 5, and axes in ring 6) were for analysis as these parameters generate the best SRRF images. Then these generated images are the super-resolved images having one pixel split into 25 pixels (5×5 pixels) creating a high-resolution image. To maintain the uniformity in reconstructed images background of the artifacts was removed at the same parameter. For scaling, color coding merging, all the images were processed to get the super-resolved final version by implementing ImageJ software. The quantitative and statistical analysis was performed with the help of the custom-made scripts in MATLAB and ImageJ macro language.

#### d. Image Processing and Analysis

ImageJ (National Institutes of Health): This open-source image processing software was used for analyzing fluorescence intensity and colocalization. ImageJ offers various tools and plugins that facilitate the quantification of fluorescence signals and the assessment of colocalization between different fluorescent markers.

### 11. Statistical Analysis

Statistical analyses were carried out using SPSS Statistics software. For each experimental condition, a sample size of 10 was utilized. The data were subjected to independent paired t-tests to assess the significance of observed differences between treatment groups. A threshold of p < 0.05 was chosen to determine significant differences across all analyses in the study. This criterion ensures robust evaluation of the experimental outcomes and provides confidence in the reliability of the observed effects under different experimental conditions.

## Data and Materials Availability

Most of the data we provided in the main manuscript are available with the manuscript itself and its supplementary information.

## Acknowledgements

The authors thank the Advanced Materials Research Centre (AMRC) and the Indian Knowledge System and Mental Health Applications (IKSMHA) Centre of the Indian Institute of Technology Mandi (IIT Mandi) for providing the facilities and the sophisticated instruments. All the contributing authors thank the Ministry of Education (MoE), India, for the research scholarship. CKN, SC and BM thank the IKSMHA Centre for the financial support.

## Author Contributions

CKN designed and conceptualized experiments with the help of LB, SC and BM. SC optimized the protocol and performed all the plant experiments. AS helped in imaging and result analysis. FA helped in immunostaining and super-resolution microscopy. CKN wrote the manuscript with the help of LB and SC. CKN and LB guided the complete project.

